# LIGER2: Scalable Single-Cell Integration with On-Disk Datasets

**DOI:** 10.64898/2026.09.08.750130

**Authors:** Yichen Wang, Andrew Robbins, Gaurav Gadhvi, Joshua D Welch

## Abstract

Correcting batch effects and integrating single-cell sequencing datasets has been a crucial step in large-scale biological studies. Many methods have been published for this task, with complementary strengths in various scenarios. Our previous work, LIGER, leveraging integrative non-negative matrix factorization (iNMF), stands out in providing an interpretable low-dimensional representation. To adapt to the modern need for integrating millions of cells, we developed a highly-optimized parallel factorization solution with on-demand loading from disk. The upgraded LIGER algorithm shows significant improvements in time and memory efficiency for single-cell data integration. We also developed a new downstream embedding alignment method significantly improved performance in conserving biological variation while still aligning corresponding cell types across datasets.

## INTRODUCTION

Single-cell sequencing has been proven to be an invaluable tool in biological studies for years, allowing novel insights about differences among cells within heterogeneous populations [1][2]. Numerous consortia, such as the Human Cell Atlas [3] and BRAIN Initiative, as well as efforts from individual labs [4][5][6][7][8], have contributed to rapidly scaling data generation. The integration of data from multiple experiments, however, remains a challenge. Systematic technical variation, such as batch effects, can obscure true biological variance and lead to spurious conclusions [9]. Correction of batch effects and integration of datasets is thus a prerequisite for proceeding to reliable biological inference from multi-dataset studies and atlas construction efforts.

Various computational methods have been developed to address the challenge, including Harmony [10], Scanorama [11], scVI (single-cell variational inference) [12], Seurat CCA (canonical correlation analysis) [13] and RPCA (reciprocal principal component analysis, reciprocal PCA) [14]. These popular tools have demonstrated excellent performance in removing batch effects and conserving biological variance according to benchmark studies [15].

Our previous work, LIGER (Linked Inference of Genomic Experimental Relationships), introduced an integrative non-negative matrix factorization (iNMF) framework to find metagene loading in cells, shared gene loading in metagenes, and dataset-specific gene loading in metagenes. A metagene can be interpreted as a biological program representing cell identity or activity [16], and thus guides the alignment of populations across batches. We used quantile normalization to align the cell metagene loading [17][18]. LIGER has demonstrated outstanding performance in single-cell integration and is a widely used tool that provides an interpretable low-dimensional output. However, our previous implementations still leave much to be desired in terms of scalability. Furthermore, quantile normalization can sometimes overcorrect, reducing the conservation of true biological variation during data integration.

As atlas-scale single-cell datasets become routine, efficiency is no longer a matter of convenience. In practice, integration often needs to be repeated multiple times as datasets are updated, reprocessed, or subset for targeted analyses. Methods that demand long runtime or specialized hardware impose real constraints on iteration speed, slow down downstream analyses, and limit exploratory work across alternative annotations or analysis strategies.

To address these challenges, we have developed a new parallelized factorization framework for LIGER, enabling the integration of a million cells in under ten minutes with minimal memory usage on standard hardware available to most people. We also introduce a novel downstream embedding alignment method, centroid alignment, which improves the conservation of biological variance from the original datasets while minimizing the compromise of batch effect removal. Benchmarking across diverse real atlases demonstrates that our upgraded LIGER achieves both computationally efficient single-cell integration while achieving state-of-the-art results in both biological conservation and batch correction.

## METHOD OVERVIEW

### Scalable parallelized computation

We refactored the iNMF algorithm and its variants [19][20] entirely in C++ by incorporating a published parallel NMF framework, PLANC [21] (see Supplementary Methods). To enable portability in various high-level programming environments, we released the new implementation as a standalone C++ library, libplanc, together with interface package RcppPlanc and pyplanc for R and Python users, respectively. The upgraded iNMF runs in a fully parallelized manner, processing only a chunk of data at a time per thread. This chunk-wise strategy can be applied to data already loaded into memory or to on-disk datasets (see Supplementary Methods), allowing integration of datasets that cannot be entirely loaded in RAM at once (Fig. 1A). The refactored iNMF algorithms yields results numerically identical to the previous implementations.

**Figure 1.**
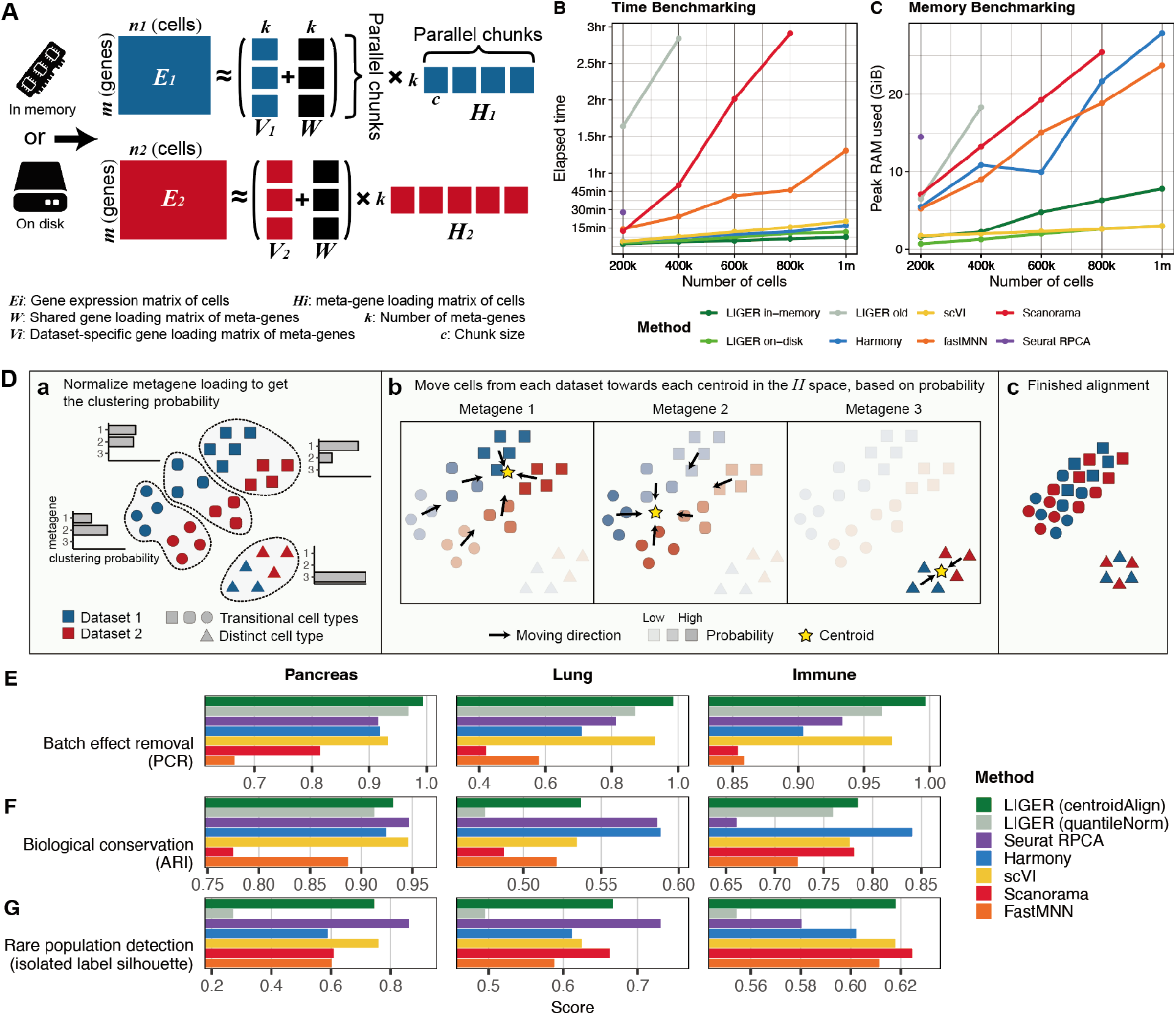
Upgraded LIGER schematics diagram and performance. **A**. The schematic diagram showing parallelized iNMF. **B**. Runtime benchmarking result comparing LIGER against other methods. **C**. Peak memory usage benchmarking result comparing LIGER against other methods. **D**. Illustrative figure of centroid alignment. The scattered points illustrate the neighboring relationship among all cells as if the dimensionality of the metagene loading matrix is reduced to 2D. a. Before aligning. b. Solving the movement in the dimension of each metagene. Arrows in the sub-panels indicate the direction of movement. c. After aligning. **E** Principal component regression (PCR) score of each method across three datasets. **F** Adjusted rand index (ARI) score of each method across three datasets. **G** F1 score of each method across three datasets, measuring the detection of isolated labels.

### Centroid alignment

We developed a new downstream cell metagene loading alignment method, centroid alignment, which serves as an optional replacement of quantile normalization in the LIGER integration workflow. In our previous work, we first assigned cluster label based on the maximum metagene loading in each cell, based on the nature of nonnegative matrix factorization (NMF) method [16]. We then applied quantile normalization on the cell metagene loading matrix to match the loading distribution of metagenes of cells belonging to the same cluster but different datasets [18]. In centroid alignment, we normalize all the metagene loading of each cell to represent the probability that a cell belongs to each cluster, which can be viewed as a type of soft clustering. This is to avoid mischaracterization of cells with multiple highly loaded metagenes and to preserve the potential trajectory of cells in a transitional state (Fig. 1Da). We next linearly transform the embedding space to align the centroids of each cluster from all datasets, weighted by soft clustering probability (Fig. 1Db). In this way, the original metagene space of each cluster from different datasets is preserved (Fig. 1Dc), rather than being distorted by enforcing distribution alignment (see Supplementary Methods).

## BENCHMARKING

### Computational performance

To assess the computational performance of the upgraded iNMF implementation, we measured its runtime, peak memory usage and scaling behavior across scRNAseq dataset ranging from the sizes of 200,000 to one million cells, in both in-memory and on-disk data modes. We used a publicly available single-cell atlas of the human hematopoietic system, where we took a subset of 1,033,203 cells from 44 batches. We further randomly subset it to a series of desired test sizes. We compared the measurement of the equivalent steps from other widely used methods, including Harmony [10], Scanorama [11], scVI [12], Seurat RPCA [14] and fastMNN [22]. We set a time limit for each test to the maximum of 3 hours and a memory limit to the maximum of 320 GiB. We allowed 32 cores for all methods equipped with multi-core processing. All methods were run on the same hardware to ensure comparability, except that scVI was run with an additional GPU as it does not scale to the tested dataset sizes without a GPU (see Supplementary Methods).

Across all dataset sizes tested, the upgraded in-memory iNMF consistently achieved the fastest runtime. The on-disk version, with little compromise in runtime, required the lowest memory usage. The on-disk iNMF ran still faster than any other method. The in-memory iNMF integrated a million cells in under 8 minutes with using 8 GiB peak memory, and the on-disk iNMF completed the same task in under 12 minutes with using only 3 GiB peak memory (Fig. 1B, C). In contrast, the previous version of LIGER, Seurat RPCA and scanorama failed to complete the integration of a million cells within the given time and memory limit. While Harmony shows competitive runtime performance, it required the most memory usage among the methods that completed the integration of a million cells. scVI, with additional GPU support enabled, was able to show competitive runtime and memory performance. The runtime and memory usage of Harmony, scVI, and LIGER all scaled nearly linearly with the number of cells, whereas fastMNN exhibited superlinear growth at larger cell counts. In addition to cell count, we observed near-linear scaling of runtime and memory usage with the number of highly variable genes (HVGs) selected, the number of metagenes targeted, and the number of batches (Supplementary Fig. S1).

### Integration quality

We evaluated the integration quality of centroid-aligned iNMF embedding using widely accepted single-cell integration benchmarking metrics. These metrics are divided into two categories. For evaluating the removal of batch effect, we tested the graph connectivity, batch-level local inverse Simpson’s index (LISI) (iLISI)[10], k-nearest neighbour batch effect test (kBET)[23], batch-level principal component regression (PCR)[23], and batch-level average silhouette width (ASW)[23]. For evaluating the conservation of original biological variation, referred to as bio-conservation later, we used normalized mutual information (NMI)[24], adjusted rand index (ARI)[25], label-level ASW, label-level LISI (cLISI). Additionally, we assessed the ability of rare population identification using isolated label F1 score, isolated label silhouette score[15]. The benchmarking was performed on three publicly available datasets of various complexities. Each of the three datasets was constructed by collecting related batches from multiple studies and therefore inherits distinctive batch effects. These datasets have been widely used to assess integration methods and were previously curated for quality control and annotation [15]. To demonstrate the improvement in the LIGER integration workflow, we benchmarked the old workflow, which uses quantile normalization to align the metagene loading produced by iNMF. Meanwhile, we also benchmark the other integration methods previously mentioned using their latest versions (see Supplementary Methods).

The centroid-aligned iNMF embedding effectively removes batch effects from the datasets benchmarked (Fig. 1E, Supplementary Fig. S2). Out of all the metrics that measure the effectiveness of batch removal, we show that our new method consistently achieves the highest score in the PCR test. We do not expect a consistent improvement in the batch removal scores as compared to the quantile normalization method, as the latter is prone to over-correction and can produce misleadingly high batch removal scores. Indeed, quantile normalization gets higher scores in the iLISI tests across all datasets, kBET tests in the lung and immune datasets, and batch-level ASW tests in the pancreas and lung datasets.

The use of centroid alignment on the iNMF result improves the conservation of original biological variation compared to using quantile normalization (Fig. 1F,G, Supplementary Fig. S2). We show that centroid alignment gets higher bio-conservation scores than quantile normalization in the test of NMI, ARI and cLISI in all datasets, as well as cell-cycle conservation score in the pancreas and immune datasets. In comparison with other methods, LIGER shows competitive capability in bio-conservation. There is no one method that consistently gets the highest score in a metric across all datasets, and each method can receive a low score in a metric-dataset pair.

Notably, centroid alignment largely improves the capability of detecting rare cell types that are only partially contributed by batches in an integration task (Fig. 1G, Supplementary Fig. S3). Centroid-aligned iNMF embedding consistently achieves improved scores compared to the quantile-normalized embedding, and gets the highest isolated label F1 score in the pancreas dataset.

Besides the quantitative benchmarking, we also demonstrate that centroid alignment is capable of preserving a global transitional trajectory on a 2D visualization embedding, using uniform manifold approximation and projection (UMAP) [26] (Supplementary Fig. S5). The immune dataset was intentionally collected to include both peripheral blood and bone marrow samples and therefore contains hematopoietic stem and progenitor cells (HSPCs) and progenitors of different lineages. In the classical hematopoietic hierarchy, HSPCs give rise to common myeloid progenitors (CMPs) and common lymphoid progenitors (CLPs) through an intermediate multipotent progenitor (MPP) stage. CMPs further differentiate into megakaryocyte–erythroid progenitors (MEPs) and granulocyte–macrophage progenitors (GMPs). MEPs generate erythrocytes and megakaryocytes, whereas GMPs give rise to granulocytes and monocytes [27]. From the UMAP generated from centroid-aligned iNMF embedding, we can clearly observe a continuous branch from HSPCs to monocyte progenitors, monocytes, and eventually monocyte-derived dendritic cells (moDCs). On the other side of the HSPC cluster, we can see another continuous branch through erythroid progenitors and megakaryocyte progenitors, and it ends at erythrocytes. The structure agrees with the prior knowledge. A similar structure is only observed from Seurat RPCA integration, whereas the other methods produce breakage on the trajectory (quantile normalization, Harmony, scVI, and Scanorama), overlap with unrelated cell types (scVI), and twisted structure (fastMNN).

## SOFTWARE AND DATA AVAILABILITY

The LIGER method described in this work is presented in the R package rliger (version 2.2.0), published on CRAN (https://cran.r-project.org/package=rliger) and publicly available on GitHub (https://github.com/welch-lab/liger). The RcppPLANC package we developed to implement parallel iNMF is also available on CRAN and GitHub. The human immune dataset used for benchmarking computational performance is available at Human Cell Atlas (https://explore.data.humancellatlas.org/projects/cc95ff89-2e68-4a08-a234-480eca21ce79). The curated datasets used for benchmarking integration quality is available on at figshare (https://figshare.com/articles/dataset/Benchmarking_atlas-level_data_integration_in_single-cell_genomics_-_integration_task_datasets_Immune_and_pancreas_/12420968).

## ACKNOWLEDGMENTS

This work was supported by NIH grant R01HG010883 to J.D.W.

## REFERENCES

[1] Valentine Svensson, Roser Vento-Tormo, and Sarah A Teichmann. “Exponential scaling of single-cell RNA-seq in the past decade”. In: Nature Protocols 13 (Mar. 2018), pp. 599–604. DOI: 10.1038/nprot.2017.149.

[2] Mariano I. Gabitto, Kyle J. Travaglini, Victoria M. Rachleff, et al. “Integrated multimodal cell atlas of Alzheimer’s disease”. In: Nature Neuroscience 27 (Oct. 2024), pp. 2366–2383. DOI: 10.1038/s41593-024-01774-5.

[3] Aviv Regev, Sarah A Teichmann, Eric S Lander, et al. “The Human Cell Atlas”. In: eLife (2017). DOI: 10.7554/eLife.27041.

[4] Zizhen Yao, Cindy T. J. van Velthoven, Michael Kunst, et al. “A high-resolution transcriptomic and spatial atlas of cell types in the whole mouse brain”. In: Nature 624 (14 Dec. 2023). DOI: 10.1038/s41586-023-06812-z.

[5] Rasa Elmentaite, Natsuhiko Kumasaka, Kenny Roberts, et al. “Cells of the human intestinal tract mapped across space and time”. In: Nature 597 (9 Sept. 2021), pp. 250–255. DOI: 10.1038/s41586-021-03852-1.

[6] Xingfan Huang, Jana Henck, Chengxiang Qiu, et al. “Single-cell, whole-embryo phenotyping of mammalian developmental disorders”. In: Nature 623 (Nov. 2023), pp. 772–781. DOI: 10.1038/s41586-023-06548-w.

[7] Renying Wang, Peijing Zhang, Jingjing Wang, et al. “Construction of a cross-species cell landscape at single-cell level”. In: Nucleic Acids Research 51 (2 Jan. 2023), pp. 501–516. DOI: 10.1093/nar/gkac633.

[8] CZI Single-Cell Biology Program, Shibla Abdulla, Brian Aevermann, et al. “CZ CELL×GENE Discover: A single-cell data platform for scalable exploration, analysis and modeling of aggregated data”. In: bioRxiv (2023). DOI: 10.1101/2023.10.30.563174.

[9] Stephanie C Hicks, F William Townes, Mingxiang Teng, et al. “Missing data and technical variability in single-cell RNA-sequencing experiments”. In: Biostatistics 19 (4 Oct. 2018), pp. 562–578. DOI: 10.1093/biostatistics/kxx053.

[10] Ilya Korsunsky, Nghia Millard, Jean Fan, et al. “Fast, sensitive and accurate integration of single-cell data with Harmony”. In: Nature Methods 16 (12 Dec. 2019), pp. 1289–1296. ISSN: 1548-7091. DOI: 10.1038/s41592-019-0619-0.

[11] Brian L. Hie, Soochi Kim, Thomas A. Rando, et al. “Scanorama: integrating large and diverse single-cell transcriptomic datasets”. In: Nature Protocols 19 (8 Aug. 2024), pp. 2283–2297. ISSN: 1754-2189. DOI: 10.1038/s41596-024-00991-3.

[12] Romain Lopez, Jeffrey Regier, Michael B. Cole, et al. “Deep generative modeling for single-cell transcriptomics”. In: Nature Methods 15 (12 Dec. 2018), pp. 1053–1058. ISSN: 1548-7091. DOI: 10.1038/s41592-018-0229-2.

[13] Andrew Butler, Paul Hoffman, Peter Smibert, et al. “Integrating single-cell transcriptomic data across different conditions, technologies, and species”. In: Nature Biotechnology 36 (2018), pp. 411–420. DOI: 10.1038/nbt.4096.

[14] Tim Stuart, Andrew Butler, Paul Hoffman, et al. “Comprehensive Integration of Single-Cell Data”. In: Cell 177 (7 2019), 1888–1902.e21. DOI: 10.1016/j.cell.2019.05.031.

[15] Malte D. Luecken, M. Büttner, K. Chaichoompu, et al. “Benchmarking atlas-level data integration in single-cell genomics”. In: Nature Methods 19 (1 Jan. 2022), pp. 41–50. ISSN: 1548-7091. DOI: 10.1038/s41592-021-01336-8.

[16] Jean-Philippe Brunet, Pablo Tamayo, Todd R. Golub, et al. “Metagenes and molecular pattern discovery using matrix factorization”. In: Proceedings of the National Academy of Sciences 101.12 (2004), pp. 4164–4169. DOI: 10.1073/pnas.0308531101.

[17] Joshua D. Welch, Velina Kozareva, Ashley Ferreira, et al. “Single-Cell Multi-omic Integration Compares and Contrasts Features of Brain Cell Identity”. In: Cell 177 (7 June 2019), 1873–1887.e17. ISSN: 00928674. DOI: 10.1016/j.cell.2019.05.006.

[18] Jialin Liu, Chao Gao, Joshua Sodicoff, et al. “Jointly defining cell types from multiple single-cell datasets using LIGER”. In: Nature Protocols 15 (11 Nov. 2020), pp. 3632–3662. ISSN: 1754-2189. DOI: 10.1038/s41596-020-0391-8.

[19] Chao Gao, Jialin Liu, April R. Kriebel, et al. “Iterative single-cell multi-omic integration using online learning”. In: Nature Biotechnology 39 (8 Aug. 2021), pp. 1000–1007. ISSN: 1087-0156. DOI: 10.1038/s41587-021-00867-x.

[20] April R. Kriebel and Joshua D. Welch. “UINMF performs mosaic integration of single-cell multi-omic datasets using nonnegative matrix factorization”. In: Nature Communications 13 (1 Feb. 2022), p. 780. ISSN: 2041-1723. DOI: 10.1038/s41467-022-28431-4.

[21] Srinivas Eswar, Koby Hayashi, Grey Ballard, et al. “PLANC: Parallel Low-rank Approximation with Nonnegativity Constraints”. In: ACM Transactions on Mathematical Software 47.3 (June 2021). DOI: 10.1145/3432185.

[22] Feng Zhang, Yu Wu, and Weidong Tian. “A novel approach to remove the batch effect of single-cell data”. In: Cell Discovery 5.46 (2019). DOI: 10.1038/s41421-019-0114-x.

[23] Maren Büttner, Zhichao Miao, F. Alexander Wolf, et al. “A test metric for assessing single-cell RNA-seq batch correction”. In: Nature Methods 16 (2018), pp. 43–49. DOI: 10.1038/s41592-018-0254-1.

[24] Fabian Pedregosa, Gaël Varoquaux, Alexandre Gramfort, et al. “Scikit-learn: Machine Learning in Python”. In: Journal of Machine Learning Research 12 (85 2011), pp. 2825–2830.

[25] Lawrence Hubert and Phipps Arabie. “Comparing partitions”. In: Journal of Classification 2 (1985), pp. 193–218. DOI: 10.1007/BF01908075.

[26] Leland McInnes 1, John Healy, Nathaniel Saul, et al. “UMAP: Uniform Manifold Approximation and Projection”. In: The Journal of Open Source Software 3 (29 2018), p. 861. DOI: 10.21105/joss.00861.

[27] Stuart H. Orkin and Leonard I. Zon. “Hematopoiesis: An Evolving Paradigm for Stem Cell Biology”. In: Cell 132 (4 Feb. 2008), pp. 631–644. DOI: 10.1016/j.cell.2008.01.025.

