## Supplementary methods and figures for "LIGER2: Scalable Single-Cell Integration with On-Disk Datasets"

##### Parallelized iNMF

The integrative non-negative matrix factorization (iNMF) algorithm is previously defined as finding matrix $W$, $V$ and $H$ that minimize the objective function in eq. 1.

$$W,V,H = arg \min_{W \in \mathbb{R}_{\geq 0}^{m \times k}, \{V_d \in \mathbb{R}_{\geq 0}^{m \times k}, H_d \in \mathbb{R}_{\geq 0}^{k \times n_d}\}_{d=1}^{D}} \sum_{d=1}^{D} \|E_d - (W + V_d)H_d\|_F^2 + \lambda \sum_{d=1}^{D} \|V_d H_d\|_F^2, \quad E_d \in \mathbb{R}_{\geq 0}^{m \times n_d} \tag{1}$$

In the objective function, there are $D$ batches of dataset to be integrated, and $E_d$ is the scaled expression value of $m$ shared highly variable genes from $n_d$ cells in the $d$'th dataset. With targeting at $k$ metagenes, $W$ denotes the shared gene loading matrix of factorized metagenes, $V_d$ is the dataset-specific metagene gene loading matrix for the $d$'th dataset, and $H_d$ is the cell metagene loading matrix for the $d$'th dataset. $\lambda$ is a regularization parameter. All the matrices are non-negative. $\|.\|_F^2$ denotes Frobenius norm of a matrix.

To find the $W$, $V$ and $H$ matrices that minimize the objective error with $E$ matrices fixed, we optimize the function with an alternating nonnegative least squares (ANLS) framework. With all these matrices randomly initialized, we iteratively solve and update each type of matrix with solving non-negative least square (NNLS) problem. An NNLS problem is generalized as finding non-negative matrix $X$ that minimizes the objective function in eq. 2 when non-negative matrices $C$ and $B$ are given.

$$X = \arg \min_{X \in \mathbb{R}_{\geq 0}} \|CX - B\|_F^2, \quad C, B \in \mathbb{R}_{\geq 0} \tag{2}$$

The exact updating steps in the ANLS framework of iNMF solves $H_d$, $V_d$ and $W$ in the order for each iteration using eq. 3, 4, 5.

$$H_d = \arg \min_{H_d \in \mathbb{R}_{\geq 0}^{k \times n_d}} \left\| \begin{pmatrix} W + V_d \\ \sqrt{\lambda} V_d \end{pmatrix} H_d - \begin{pmatrix} E_d \\ \mathbf{0}_{m \times n_d} \end{pmatrix} \right\|_F^2 \tag{3}$$

$$V_d = \arg \min_{V_d \in \mathbb{R}_{\geq 0}^{m \times k}} \left\| \begin{pmatrix} H_d^\intercal \\ \sqrt{\lambda} H_d^\intercal \end{pmatrix} V_d^\intercal - \begin{pmatrix} E_d^\intercal - H_d^\intercal W^\intercal \\ \mathbf{0}_{n_d \times m} \end{pmatrix} \right\|_F^2 \tag{4}$$

$$W = \arg \min_{W \in \mathbb{R}_{\geq 0}^{m \times k}} \left\| \begin{pmatrix} H_1^\intercal \\ \vdots \\ H_d^\intercal \end{pmatrix} W^\intercal - \begin{pmatrix} E_1^\intercal - H_1^\intercal V_1^\intercal \\ \vdots \\ E_d^\intercal - H_d^\intercal V_d^\intercal \end{pmatrix} \right\|_F^2 \tag{5}$$

To parallelize the solution of each column in $X$ of eq. 2, we introduced block principal pivoting method (BPPNNLS) [1], frameworked in C++ library PLANC[2], which allows for individual columns of $X$ and $B$ to be used independently. Besides, BPPNNLS requires the input of $C^\intercal C$ and $C^\intercal B$, which can be derived from original iNMF elements. The resulting $C^\intercal C$ matrix is always of $k$ by $k$ dimensionality. The dimensionality of $C^\intercal B$ is $k$ by $n$ when solving $H_d$ and $k$ by $m$ when solving $V_d$ and $W$. As the practical value of $k$ is often around 20 to 50, the total memory footprint is largely reduced. In practice, each column of $B$, denoted as $b$, is used in each parallel worker (eq. 6- 11). Let $i,.$ denotes a row vector of a matrix and $.,j$ denotes a column vector of a matrix,

For solving $H_d$:

$$C^{\intercal}C = (W^{\intercal} + V_d^{\intercal})(W + V_d) + \lambda V_d^{\intercal} V_d \tag{6}$$
$$C^{\intercal}b = (W^{\intercal} + V_d^{\intercal})E_{d:.,j} \tag{7}$$

For solving $V_d$:

$$C^{\intercal}C = (1+\lambda)H_d H_d^{\intercal} \tag{8}$$
$$C^{\intercal}b = H_d E_{d:i,.}^{\intercal} - H_d H_d^{\intercal} W_{i,.}^{\intercal} \tag{9}$$

For solving $W$:

$$C^{\intercal}C = \sum_{d=1}^{D} H_d H_d^{\intercal} \tag{10}$$
$$C^{\intercal}b = \sum_{d=1}^{D} H_d E_{d:i,.}^{\intercal} - H_d H_d^{\intercal} V_{d:i,.}^{\intercal} \tag{11}$$

The parallelization is implemented with OpenMP library, supported on Windows and Linux operating systems. The linear algebra computation is accelerated with OpenBLAS library. Although OpenMP is not supported on macOS operating systems, machines equipped with modern Apple Silicon processors show competitive performance in single-thread mode.

#### On-disk iNMF

To enable fast parallel iNMF on large on-disk datasets, we construct data container objects that hold the handles for retrieving indexed data from on-disk HDF5 files. The preprocessing steps, including normalization, HVG selection, and scaling, also benefit from such a data structure. When solving each type of matrix in an ANLS iteration, each dataset is traversed in order. We load only one dataset into memory at a time, in full, and parallelize it into chunks as described above. The iterator cleans up the loaded data and closes the file handle when one dataset is temporarily done, and loads the next dataset. This allows fast overall speed as the process does not access on-disk data every time a chunk is needed, also known as a reduction in IO (input and output) overhead. Loading the scaled expression of HVGs of all cells from one dataset is feasible for the goal of minimizing memory usage because the upstream non-centering scaling strategy retains the sparsity of the expression data, allowing the data to be stored and represented in a sparse and compressed form. As of today, empirically, a technical batch of scRNAseq data often captures fewer than tens of thousands of cells. Additionally, only a few thousand HVGs are selected. Furthermore, the raw counts data or the normalized data are not loaded during the iNMF process. Overall, the memory requirement for loading the scaled data of a single batch is totally affordable by a personal computer.

#### Runtime and peak memory usage benchmarking

The dataset used for evaluating the computational performance of all methods was acquired from Human Cell Atlas, entitled A single cell immune cell atlas of human hematopoietic system, published by Broad Institute. Out of the whole atlas that contain 1.5M cells, we take the subset of only standard scRNAseq assay that consists of 1.3M cells from 44 donors. We merged the samples from the same donor as one batch and establish the batch variable being used in all tests. Subsequently, we randomly subset the selected cells to the sizes of 200K, 400K, 600K and 800K to test for the linear scalability of each method. We performed this set of benchmarking on a high-performance computing cluster (Lighthouse, University of Michigan) node equipped with an AMD EPYC 7H12 64-core processor, using a Linux operating system. We allocated 320 gibibytes (GiB) of memory and 32 cores for each task.

For a fair comparison across all the methods, we used consistent settings that primarily affect runtime speed and memory usage. We used 4,000 variable genes and 50 latent dimensions across all methods and all task sizes. We measured LIGER's performance using both the old implementation (R package version 1.0.1) and the upgraded version (R package version 2.2.0). Specifically, we measured `runINMF` and `centroidAlign` for the new version, and `optimizeALS` and `quantile_norm` for the old version. The LIGER integration steps (i.e. iNMF followed by centroid alignment or quantile normalization) are considered as reducing the variable gene expression space to a low-dimensional representation that is ready to use for community-based clustering and creating visualization. Therefore, in benchmarking the other methods, we measured the function calls that are considered to be involved in this procedure. For harmony, we measured the total time of the `RunPCA` in Seurat (R package version 5.2.1) and the function `RunHarmony` within Harmony (R package version 1.2.3). For FastMNN, we measured the function `fastMNN` within batchelor (R package version 1.20.0), with an additional merge tree specified to account for batches from the same donors first and the same tissues next. We noticed that the default merge tree formation is significantly slower than customized ones with prior knowledge. For Seurat RPCA method, we measured the total time of the function `RunPCA` and `IntegrateLayers`, with method function set to `RPCAIntegration` and an additional merge tree specified similar to what was given to FastMNN. For Scanorama, we measured `run_scanorama` in scanorama (Python package version 1.7.4). For scVI (Python package scvi-tools version 1.3.0), we measured the total time of registering the data, building the SCVI model with two hidden layers and a negative binomial distribution, and training the model. Time and peak memory usage were measured with the R package peakRAM (version 1.0.2) for R-based methods. In Python, we used the sys module for time measurement and memory_profiler (version 0.61.0) for peak memory measurement.

We also benchmark the time and peak memory usage of the upgraded LIGER methods across various settings to test for LIGER's linear scalability against different parameters. We benchmarked the number of batches being involved while having the same total number of cells. For a given number of batches to test, we randomly assigned group labels to the existing 44 batches and concatenated the raw data within the same group. We tested the number from 20 to 40 with a 5-step increment. We next benchmarked the number of variable genes selected from 2,000 to 8,000 genes with a 1,000-step increment, while fixing the number of metagenes at 40 and the number of merged batches at 20. Lastly, we benchmarked the number of metagenes from 20 to 50 with a 5-step increment, fixing the number of variable genes at 4,000 and the number of merged batches at 20.

#### Centroid alignment

To reduce residual batch-specific variation in the cell metagene loading matrices ($H_d$), we developed the new adjustment method based on weighted linear modeling. The approach assumes that batch effects can be modeled as additive offsets in the embedding space, varying across metagenes.

The method leverages the interpretability of NMF-derivative approach. We first horizontally concatenate all the cell metagene loading matrices from each dataset $H_d$ to get $H' \in \mathbb{R}_{\geq 0}^{K\times N}$ (eq. 12). $H'$ has $K$ rows for number of metagenes and $N$ columns for total number of cells. We normalize the metagene loading of each cell to a unity sum to create $P \in [0,1]^{K\times N}$, which represents the metagene membership probability (eq. 13). Meanwhile, we scale $H'$ to create $X \in \mathbb{R}^{K\times N}$, which ensures that the variance of loading is the same across metagenes (eq. 14,15). $X$ is going to be adjusted subsequently and will serve as the base of the final output. $\overline{H'_{k,.}}$ is a scalar for the mean of the $k$'th row of $H'$.

$$H' = \begin{pmatrix} H_1 & H_2 & \cdots & H_d \end{pmatrix} \tag{12}$$

$$P_{k,n} = \frac{H'_{k,n}}{\sum_{i=1}^{K} H'_{i,n}} \tag{13}$$

$$\overline{H'_{k,.}} = \frac{\sum_{j=1}^{N} H'_{k,j}}{N} \tag{14}$$

$$X_{k,n} = \frac{H'_{k,n} - \overline{H'_{k,.}}}{\sqrt{\sum_{j=1}^{N} (H'_{k,j} - \overline{H'_{k,.}})/(N-1)}} \tag{15}$$

We build a one-hot matrix $\phi \in \{0,1\}^{D\times N}$ to describe the dataset membership of each cell. We further extend $\phi$ with an intercept term to construct $\phi^{\star} \in \{0,1\}^{(D+1)\times N}$ (eq. 16).

$$\phi^{\star} = \begin{pmatrix} 1 & \cdots & 1 \\ \phi_{1,1} & \cdots & \phi_{1,N} \\ \vdots & \ddots & \vdots \\ \phi_{D,1} & \cdots & \phi_{D,N} \end{pmatrix}, \phi_{d,n} = \begin{cases} 1, & \text{if cell n belongs to dataset d,} \\ 0, & \text{otherwise.} \end{cases} \tag{16}$$

For each metagene $k$, we estimate a coefficient matrix $S_k \in \mathbb{R}^{(1+D)\times K}$ via ridge-regularized weighted least squares (eq. 17).

$$S_k = (\phi^{\star}\mathrm{diag}(P_{k,.})\phi^{\star\top} + \rho I)^{-1}\phi^{\star}\mathrm{diag}(P_{k,.})X^{\top} \tag{17}$$

Here, $\mathrm{diag}(P_{k,.}) \in [0,1]^{N\times N}$ is the diagonal matrix of weights for metagene $k$ (eq. 18). $\rho$ is a ridge penalty hyperparameter used to stabilize the estimation of the shift coefficients. Larger values of $\rho$ introduce stronger shrinkage to the batch-dependent shift terms, and thus weaken the correction power. In contrast, smaller positive values allow the model to more correct the batch-specific effects, but may increase sensitivity to noise. In practice, we adopt a default of 1 to allow for an effective correction result.

$$\mathrm{diag}(P_{k,.}) = \begin{pmatrix} P_{k,1} & \cdots & 0 \\ \vdots & \ddots & \vdots \\ 0 & \cdots & P_{k,N} \end{pmatrix} \tag{18}$$

With the coefficient estimated, we exclude the intercept term to only correct the embedding $X$ with the batch-specific variance, using $S_{k[2:(D+1),.]} \in \mathbb{R}^{D\times K}$. The batch-specific offset for each cell $n$ is subtracted from the embedding, weighted by its factor memberships (eq. 19). In this way, the centroids of cells of the same identity from different batches are moved towards each other and aligned, and cells with multiple possible identities are automatically placed at a proper intermediate position in the corrected space. $\hat{X} \in \mathbb{R}^{K\times D}$ represents the final corrected embedding.

$$\hat{X}_{.,n} = X_{.,n} - \sum_{k=1}^{K} P_{k,n} S_{k[2:(D+1),.]}^{\top} \phi^{*}_{2:(D+1),n} \tag{19}$$

#### Batch effect removal and bio-conservation benchmarking

We obtained the curated immune, lung and pancreas datasets from the shared repository of the work of scIB [3]. The curated data comes with raw counts, batch assignment and cell type annotation. They additionally contain log+1 transformed normalized expression produced with Scran pooling size factors [4]. We integrated each of the datasets using the recommended pipeline and default parameters of each method, as described in detail below. We pre-selected 4,000 highly variable genes (HVGs) for each dataset, using scIB's function `scib.pp.hvg_batch`. These HVGs are used for all integration methods when reducing the data dimensionality.

##### *LIGER, Centroid Alignment And Quantile Normalization*

We normalized the counts of each cell to a sum of unity (`normalize`) and scaled the normalized expression of each HVG to a unit variance, without centering (`scaleNotCenter`). We then performed iNMF on the scaled data targeting at 40 metagenes (`runINMF(k=40)`). The final integrated embedding is generated with centroid alignment (`centroidAlign`). Additionally, to compare with our old method, we also performed quantile normalization on the raw iNMF result (`quantileNorm`). We used R package rliger version 2.2.0.

##### *Seurat, RPCA*

We constructed Seurat object for each batch at first. For each batch, we scaled the normalized expression of HVGs (`ScaleData`) and performed principal component analysis (PCA) (`RunPCA`). We next collected all the preprocessed objects and found the integration anchors with RPCA method (`FindIntegrationAnchors(reduction="rpca")`. After integrating the data with anchors found (`IntegrateData`), we performed the last round of scaling and PCA dimensionality reduction to obtain the final integrated low-dimensional embedding. We used R package Seurat version 5.2.1.

##### *Harmony*

We used Seurat container object to hold the datasets and perform preprocessing. We obtained log+1 transformed normalized data (`NormalizeData`), scaled the HVGs (ScaleData) within each batch, and reduced the dimensionality with all batches together using PCA (`RunPCA`). Harmony method is subsequently applied to the PCA space (`RunHarmony`) to obtain the integrated embedding. We used R package harmony version 1.2.3.

##### *scVI*

We registered the raw counts to the scVI tool (`scvi.model.SCVI.setup_anndata(layer='counts')`. We constructed the SCVI model using 2 layers and 30 latent dimensions, fitting the gene likelihood with a negative binomial distribution (`scvi.model.SCVI(n_layers=2, n_latent=30, gene_likelihood='nb')`). The latent embedding after training the model was then taken as the integrated embedding. We use Python package scvi-tools 1.3.0.

##### *Scanorama*

We performed scanorama directly on the provided HVG normalized expression using 50 dimensions (`scanorama.integrate_scanpy` The resulting low-dimensional representation from each batch were concatenated to produce the final integrated embedding. We use Python package scanorama version 1.7.4.

##### *FastMNN*

We performed FastMNN directly on the provided HVG normalized expression (`fastMNN`). The resulting low-dimensional representation were taken as the final integrated embedding. We used R package batchelor 1.20.0.

##### *Benchmarking Metrics*

With integration results of all datasets from all methods collected, we performed the pre-constructed scIB pipeline to calculate all benchmarking metrics (`scib.metrics.metrics`). We used Python package scib version 1.1.7.

#### ABBREVIATIONS

**scRNAseq** Single-cell RNA sequencing

**LIGER** Linked Inference of Genomic Experimental Relationships

**scVI** Single-cell Variational Inference

**RPCA** Reciprocal principal component analysis

**GPU** Graphics processing unit

**HVG** Highly variable genes

**iNMF** Integrative non-negative matrix factorization

**ANLS** Alternating non-negative least squares

**NNLS** Non-negative least squares

**BPPNNLS** block principal pivoting non-negative least squares

**PLANC** Parallel Low-rank Approximation with Nonnegativity Constraints

**IO** Input and output

#### REFERENCES

[1] Jingu Kim and Haesun Park. "Toward Faster Nonnegative Matrix Factorization: A New Algorithm and Comparisons". In: *2008 Eighth IEEE International Conference on Data Mining*. IEEE, Dec. 2008, pp. 353–362. ISBN: 978-0-7695-3502-9. DOI: 10.1109/ICDM.2008.149.

[2] Srinivas Eswar, Koby Hayashi, Grey Ballard, et al. "PLANC: Parallel Low-rank Approximation with Nonnegativity Constraints". In: *ACM Transactions on Mathematical Software* 47.3 (June 2021). DOI: 10.1145/3432185.

[3] Malte D. Luecken, M. Büttner, K. Chaichoompu, et al. "Benchmarking atlas-level data integration in single-cell genomics". In: *Nature Methods* 19 (1 Jan. 2022), pp. 41–50. ISSN: 1548-7091. DOI: 10.1038/s41592-021-01336-8.

[4] Lun ATL, McCarthy DJ, and Marioni JC. "A step-by-step workflow for low-level analysis of single-cell RNA-seq data with Bioconductor [version 2; peer review: 3 approved, 2 approved with reservations]". In: *F1000Research* (2016), p. 2122.

### SUPPLEMENTARY FIGURES

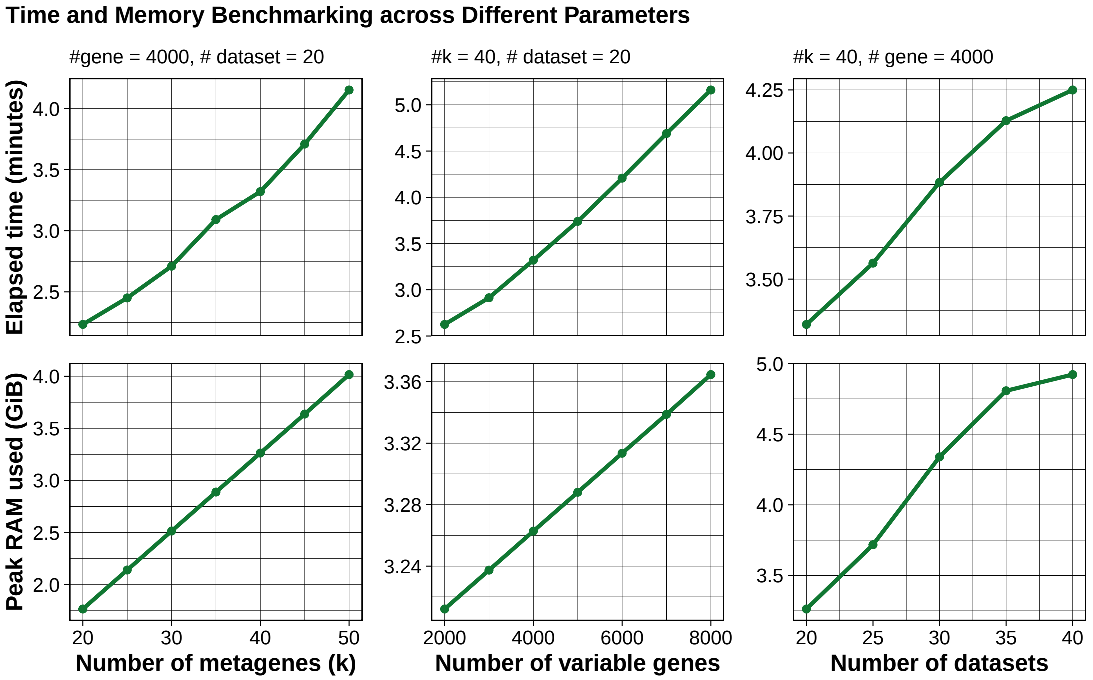


**Supplementary Figure 1.** Benchmarking runtime and memory scalability across different factors in iNMF. The first row demonstrates the runtime benchmarking and the second row demonstrates the peak memory usage. The test in the first column controls the number of variable genes and datasets being integrated and tests iNMF across different number of metagenes being factorized into. The test in the second column controls the number of metagenes and datasets and tests across different numbers of variable genes used. The test in the third column controls the number of metagenes and variable genes and tests across different number of datasets being integrated.

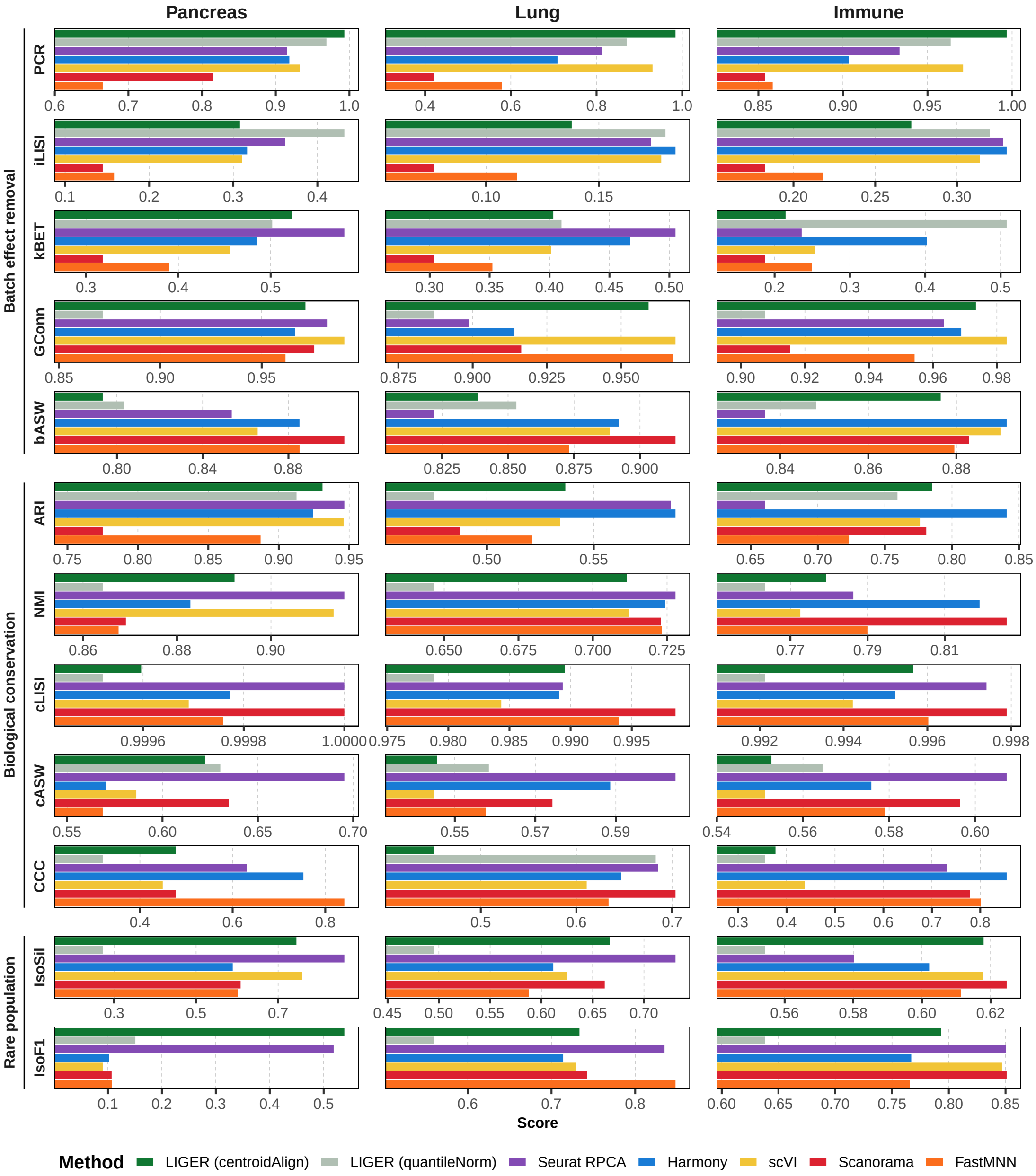


**Supplementary Figure 2.** Scores of batch effect removal and bio-conservation benchmarking across different datasets and methods. Each panel shows the score of one benchmarking metric of all methods focusing on one dataset. **PCR**: principal component regression. **iLISI**: batch-level LISI (local inverse Simpson's index) score. **kBET**: k-nearest neighbour batch effect test. **GConn**: graph connectivity. **bASW**: batch-level ASW (average silhouette width). **ARI**: adjusted rand index. **NMI**: normalized mutual information. **cLISI**: cell-type-level LISI. **cASW**: cell-type-level ASW. **CCC**: cell-cycle conservation. **IsoSil**: isolated label silhouette. **IsoF1**: isolated label F1 score.

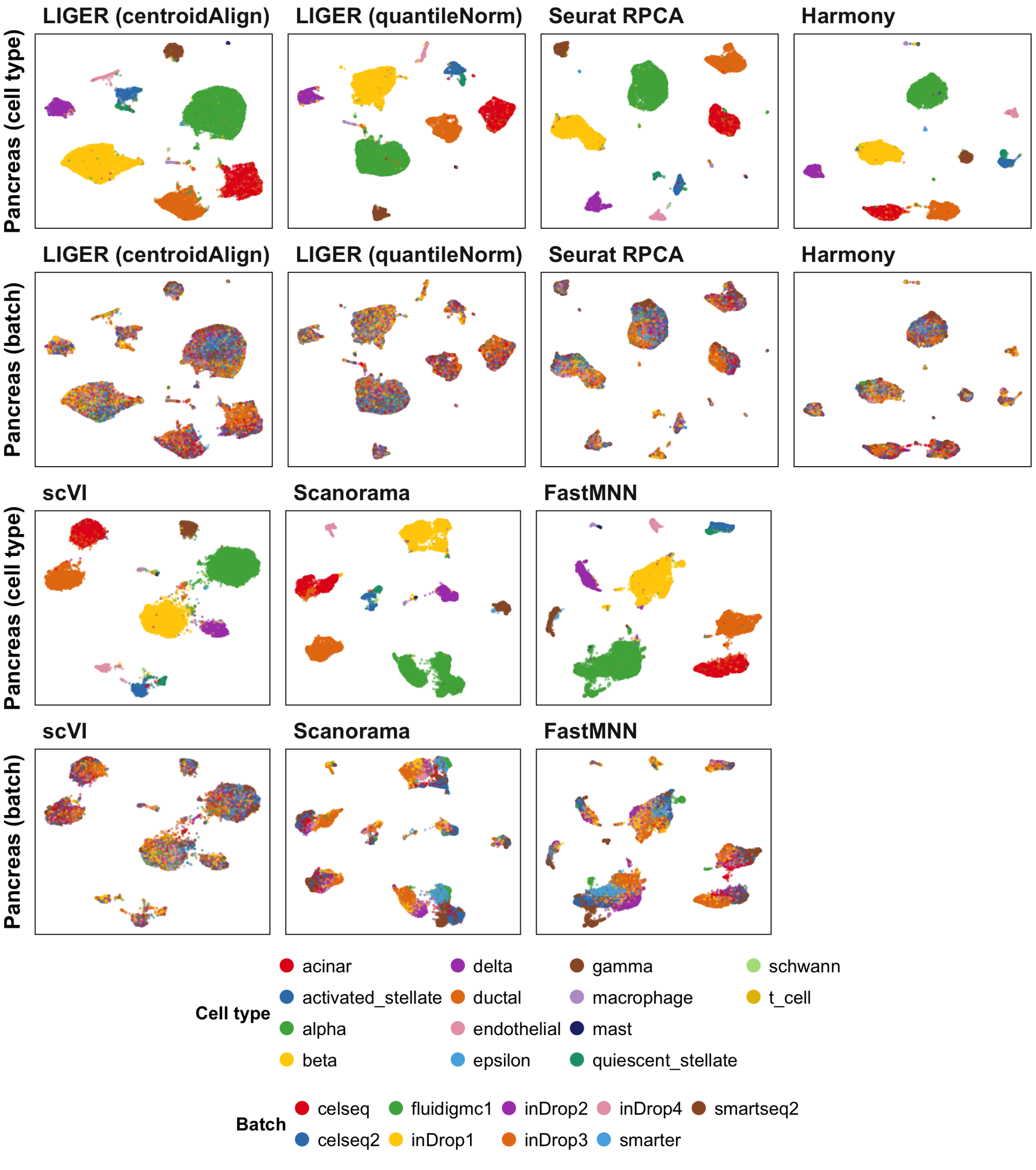


**Supplementary Figure 3.** UMAP of integration result of the pancreas dataset using different methods, with cells colored by either cell type annotation or batch belonging.

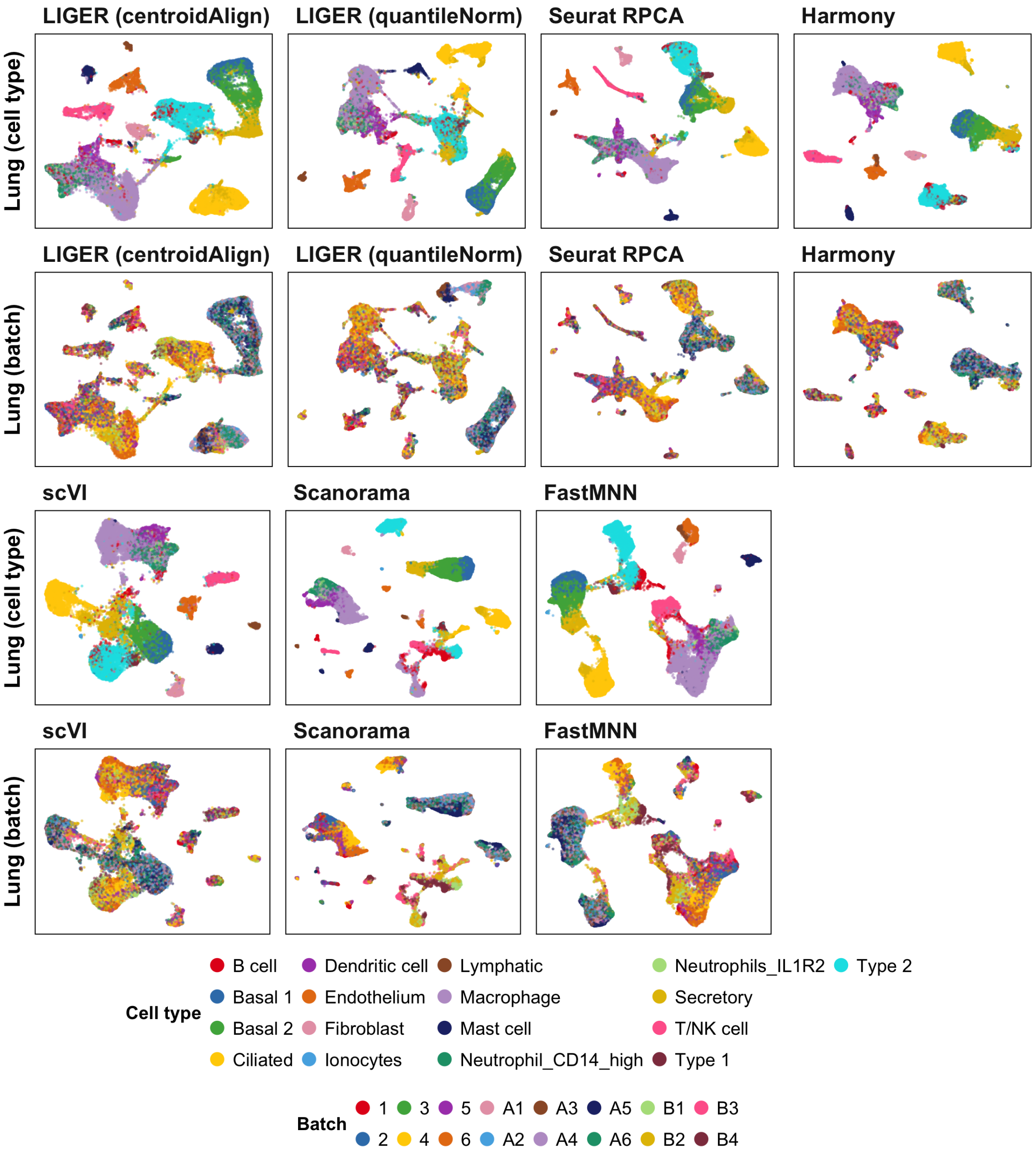


**Supplementary Figure 4.** UMAP of integration result of the lung dataset using different methods, with cells colored by either cell type annotation or batch belonging.

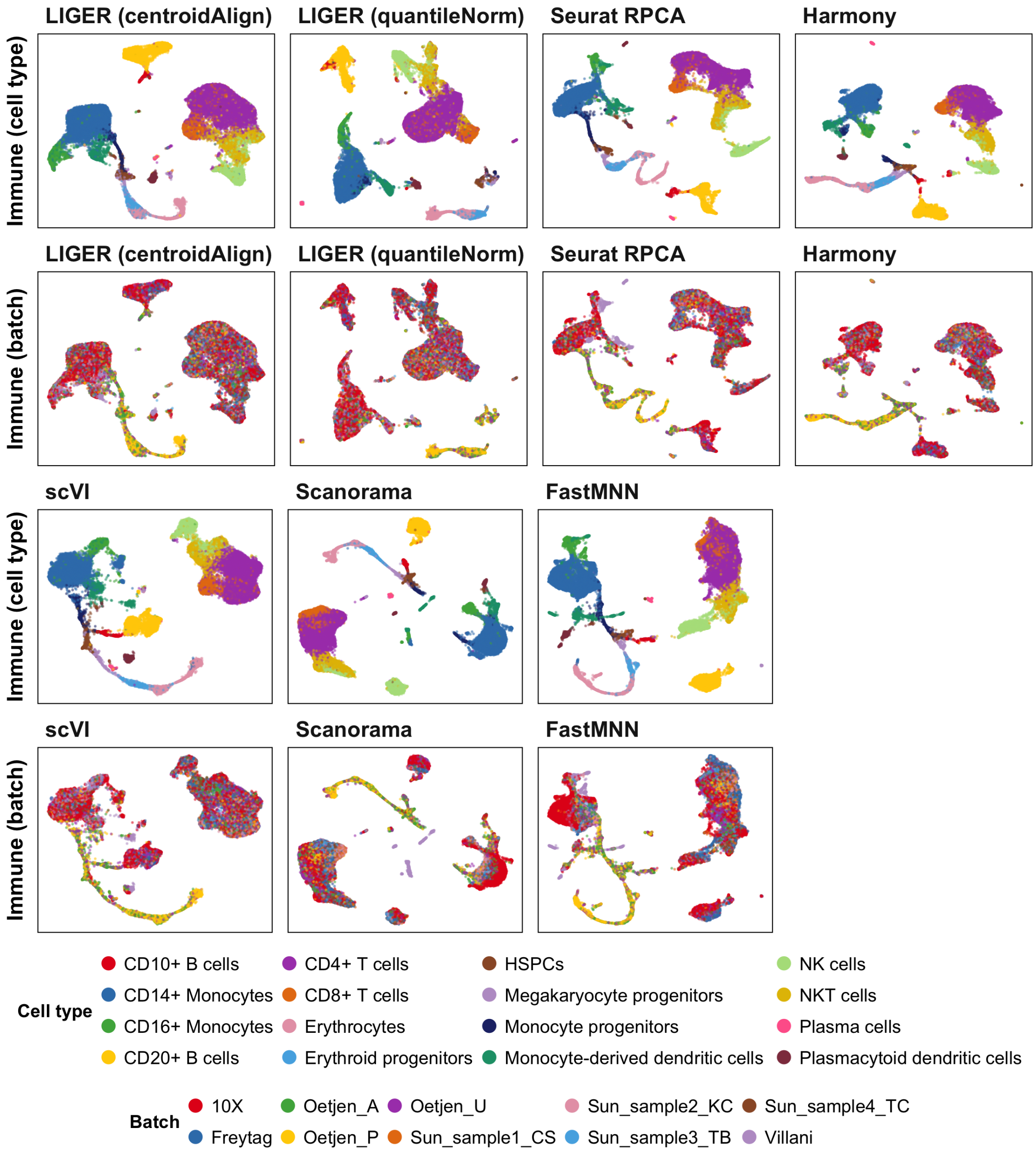


**Supplementary Figure 5.** UMAP of integration result of the immune dataset using different methods, with cells colored by either cell type annotation or batch belonging.
